# *In vitro* effects of selected angiogenesis-promoting drugs on human Ea.hy926 endothelial cells treated with critical-stage dengue hemorrhagic fever serum

**DOI:** 10.1101/2025.09.19.677322

**Authors:** Nipuni Chandrasiri, Maheshi Mapalagamage, Hansani Gunasekara, Darshan De Silva, Gayani Premawansa, Shiroma Handunnetti, Sunil Premawansa

## Abstract

Endothelial dysfunction is the major pathophysiological effect in severe dengue infection. The effects of three angiogenesis-promoting drugs, Simvastatin, Phenytoin, and Ascorbic acid on endothelial dysfunction was assessed using the simulation model using Ea.hy.926 cells treating with severe dengue hemorrhagic fever serum. Sera of patients with dengue fever (DF, n=5), dengue hemorrhagic fever (DHF, n=5), and healthy controls (HC, n=5) were treated on Ea.hy926 endothelial cells and distances were measured after 3-hour incubation. DHF-treated cells showed a significant increase in average cell-cell distances compared to HC and DF, simulating endothelial destabilization caused during DHF condition (p<0.05). Significant reductions in cell-cell distances were observed in the cells treated with 0.01 µM and 1 µM Simvastatin along with DHF serum, compared to the cells treated only with DHF serum (p<0.05). Among these two conditions tested, 0.01 µM Simvastatin showed a significant reduction in the cell-cell distances (p<0.05) and a significant increase in cell sizes (p<0.05) in the presence of DHF serum. Ascorbic acid (40 µM, 80 µM) showed no significant effect towards the endothelial monolayer destabilization in the presence of DHF serum. Both concentrations of phenytoin (10 µM, 30 µM) showed a significant increase in the average cell-cell distances compared to cells treated with DHF (p<0.05). This preliminary study suggests that low-dose Simvastatin promotes endothelial membrane formation in the presence of DHF serum and the reduction in cell-to-cell distances may not due to cell proliferation but could mainly be due to the increase in the average cell size, a known effect of Simvastatin.

## Introduction

Dengue is a mosquito-borne arboviral infectious disease mainly found in the tropical and subtropical regions of the world which causes up to 360 million symptomatic and 96 million asymptomatic infections annually (1). The global incidence of dengue has dramatically increased within the recent decades, affecting nearly half of the world’s population with an estimated 50-100 million infections, 500,000 severe dengue hemorrhagic fever (DHF) cases, and 22,000 deaths occurring each year and more than 2.5 billion people being at risk of infection. It has become a notifiable mosquito-borne disease in Sri Lanka since 1996 (2) In 2017, Sri Lanka experienced the largest dengue outbreak in the last three decades with a total of 185,688 dengue cases reported to the Epidemiology Unit of the Ministry of Health of Sri Lanka from the start of the year till the end of December. The main reason behind the escalated morbidity and mortality in the 2017 outbreak was determined to be a change in the dengue serotype. The outbreak was predominantly caused by Dengue Virus (DENV) type 2 which was not the usual type that prevailed in Sri Lanka (3).

In severe dengue infections, the critical phase/leakage phase is indicated by the onset of the plasma leakage. The complications in coagulation and vascular permeability in DHF status could be a combinatorial result of the complex interplay between the virus, host genetic makeup and host immune factors, such as viral replication, increased cell death due to infection or cytotoxic immune cells or antibodies, complement activation and increased levels of inflammatory mediators by infected or immune cells, resulting in plasma leakage (4).

>>Plasma leakage in DHF is selective and transient and usually occurs for 24-48 hours. Plasma leakage generally occurs selectively into the peritoneal and pleural spaces. The leaking normally starts slowly and increases gradually. Leaking of a sufficient volume of plasma into the interstitial cavities can result in hypovolemic shock which is short in its course, however, it can be life-threatening (4). The pathogenic mechanisms of severe dengue caused by DENV remain unresolved. However, several hypotheses have been brought forward in an effort to explain the disease severity such as DENV tropism, viral virulence, antibody-dependent enhancement (ADE), activation of the complement system and cross-reactive T cell response, host genetic factors, and cytokines (5).

An increase in vascular permeability results in plasma leakage, which is the characteristic feature of DHF, and could ultimately lead to shock syndrome, if not managed properly. The large amounts of pro-inflammatory cytokines and chemokines produced by the DENV-infected monocytes and dendritic cells, together with the activated T cells lead to dysfunction of the endothelium resulting in vascular permeability (4). The Angiopoietin-Tie-2 receptor on endothelial cells (Ang-Tie2) is an important signaling pathway involved in the remodeling and maturation of developing vasculature. The Ang-Tie2 pathway and vascular endothelial growth factor (VEGF) play an important role in the regulation of endothelial cell functions. Angiogenic factors, which are the key regulators of vascular integrity, also play important roles in the pathogenesis of severe dengue. For instance, angiopoietin-1 (Ang-1) functions to maintain vascular integrity while endothelium-derived angiopoietin-2 (Ang-2) promotes vascular leakage (6). Previous studies showed that DHF/dengue shock syndrome (DSS) patients had reduced angiopoietin-1 and increased angiopoietin-2 levels at the critical stage compared to the levels in healthy control (6, 7). This imbalance in the Ang-1:Ang-2 ratio may contribute to the transient plasma leakage in DHF/DSS. Metalloproteases also have been reported to regulate angiopoietin signaling through the cleavage of the ectodomain of the Tie-1 receptor (8–10).

Studies have shown that capillary permeability is a result of endothelial cell dysfunction, and not of endothelial cell injury, vascular leakage is known to be reversible (11). Based on this, the proposed study was designed to investigate the effects of three commercially available drugs, Simvastatin, Phenytoin, and Ascorbic acid, which are known to have angiogenic properties, on human endothelial cells treated with critical-stage DHF serum.

Simvastatin, a member of the class of statin drugs, is a 3-hydroxy3-methylglutaryl coenzyme A (HMG-CoA) reductase inhibitor that has been used to reduce a patient’s cholesterol levels. Simvastatin, in addition to its lipid-lowering properties, shows cholesterol-independent/pleiotropic benefits such as pro-angiogenic effects (12). Several studies have demonstrated the key role played by statins on endothelial cell function. Simvastatin was used in a study conducted to observe the effects of the drug on retinal microvascular endothelial cells (RMEC) and its implications for ischemic retinopathies (13). The results indicated a biphasic dose-related action of Simvastatin on RMEC where low concentrations of the drug cause a pro-angiogenic effect while higher concentrations of the drug result in anti-angiogenic effects. The results obtained were confirmed in studies conducted using human umbilical vascular endothelial cells (HUVECs) even though the two cell lines were different from each other. Concentrations lower than 5 µM were found to induce angiogenesis with 0.1 µM increasing cell proliferation and 0.01 µM promoting cell migration, sprouting, and tubulogenesis. Concentrations greater than 5 µM showed opposite effects where the cell proliferation, migration, sprouting, and tubulogenesis were inhibited. Furthermore, concentrations higher than 1 µM were found to induce cell death (13). Another study conducted by Alejandro Chade and the group in 2006 focused on the role of Simvastatin in promoting angiogenesis and prevention of microvascular remodeling in chronic renal ischemia. They tested the hypothesis that statins (Simvastatin) decrease renal injury in renal artery stenosis (RAS) by restoring angiogenesis and have found that chronic Simvastatin supplementation promoted renal angiogenesis *in vivo* and consequently restored renal hemodynamics and function (14). In a previous study, it was demonstrated that at low doses, Simvastatin shows a pro-angiogenic effect which is believed to be mediated partly through the activation of the Akt pathway as seen in mice with myocardial or hind-limb ischemia. The study also demonstrated a role for TGF-β in modulating angiogenesis by the regulation of VEGF (14). Since Simvastatin was recruited to the study as a pro-angiogenic drug, and angiogenesis is a process that is involved in cancer pathology as well, one might wonder about the implications of Simvastatin on cancer patients. Statins, however, have been rediscovered for exhibiting anticancer properties. It was found that Statins demonstrate antitumor effects such as impairment in proliferation and migration, by influencing oxidative stress-related and inflammatory tumorigenesis in vitro, in vivo experiments, and cohort studies (15–17).

Phenytoin is a commonly used non-sedative anti-epileptic agent with anticonvulsant activity (18). Phenytoin has a range of pharmacological effects in addition to its anti-epileptic activity such as inducing fibroblasts to produce increased levels of collagen and glycosaminoglycans, healing of pressure sores, venous stasis, diabetic ulcers, traumatic wounds, and also burns (19–21). The drug is found to have a secondary therapeutic role in promoting wound healing accompanying gingival hyperplasia which is characterized by increased accumulation of extracellular matrix components in gingival connective tissues (22). In a study conducted by Turan et al. in 2004, it was found that the production of VEGF increased after treating wounds with phenytoin (23). Many other studies by Pitiakoudis et al. in 2004 and DaCosta et al. in 1998 also demonstrated a role for Phenytoin in inducing angiogenesis through VEGF (24, 25). Since Phenytoin has the ability to induce angiogenesis through VEGF, it was considered as a potential drug to be used in this research to see if whether it can halt or reverse the destabilization of the endothelial cell monolayer treated with DHF serum.

Vitamin C, otherwise known as ascorbic acid, is a natural, water-soluble vitamin and a potent reducing and antioxidant agent. It is required to prevent scurvy and also has important roles in fighting bacterial infections, detoxifying reactions, and in the formation of collagen in fibrous tissue, teeth, bones, connective tissues, skin, and capillaries (26). Vitamin C enhances the endothelial synthesis and the deposition of type IV collagen to form the basement membrane of blood vessels. Recent studies have shown that the function of ascorbic acid in the endothelium extends to control endothelial proliferation and apoptosis, smooth muscle-mediated vasodilation, and endothelial permeability barrier function (26). Ascorbate (anionic form of ascorbic acid) is a key player in the process of angiogenesis as it acts as a cofactor for enzymes involved in type IV collagen synthesis (27). It stimulates the endothelial cells to proliferate by increasing the synthesis of type IV collagen which is essential for the formation of the basement membrane of the endothelial cells and also for endothelial cell adhesion. Ascorbate tightens the permeability barrier of endothelial cells and the effect is assumed to be dependent on collagen synthesis but not on collagen deposition (28). Ascorbate is necessary for the optimal function of the enzymes of collagen synthesis and many other dioxygenase enzymes (29). Such enzymes are known to regulate the transcription of proteins involved in endothelial function, proliferation, and survival. The vitamin is also considered to have a role in preventing endothelial dysfunction while systemic or localized deficiency of the vitamin is thought to be a cause for endothelial dysfunction (26). A study conducted by Daghini et al. in 2007 found that chronic administration of vitamin C with normal pigs can activate the HIF-1-VEGF pathway and lead to microvascular remodeling and cortical angiogenesis (30). Another study by (31) discovered that ascorbic acid reverses endothelial vasomotor dysfunction in patients with coronary artery disease (32). Based on such studies, vitamin C was selected for this study as an angiogenesis-promoting drug to test whether it has the possibility to prevent endothelial cell dysfunction.

The primary objective of this preliminary *in vitro* study was to investigate the potential of three angiogenesis-promoting agents—Simvastatin, Phenytoin, and Ascorbic acid—to inhibit or reverse endothelial destabilization induced by DHF serum. A secondary aim was to determine the optimal drug concentrations capable of eliciting significant morphological alterations, including changes in cell size, cell counts, and intercellular spacing, in macrovascular endothelial cells exposed to DHF serum, in comparison with sera from dengue fever patients and healthy controls.

## Materials and Methods

### Recruitment of Patients

Clinically suspected dengue patients, were recruited for the study from the Colombo North Teaching Hospital, Ragama, Sri Lanka. Informed written consent was obtained from each patient before sample collection. Ethical approval was obtained from the Ethics Review Committee, Faculty of Medicine, University of Colombo, Sri Lanka (EC-13-172, 12^th^ June 2019). NS1 and/or IgM confirmed dengue patients were further categorized as DF and DHF using clinical characterization as given in WHO, SEARO, 2011 criteria. Clinical characterization was done by the clinicians attached to wards Number 9 (male) and 12 (female) of NCTH, Ragama. An amount of 3 ml of Blood was collected from recruited and confirmed DF (n=5) and DHF (n=5) patients and sera were separated.

### Recruitment of Healthy Controls

Age and gender-matched healthy individuals (healthy controls; HC) (n=5) were recruited controls for the study. Individuals with previous dengue infection or any fever condition during the week of sample collection were excluded.

### Cell Line

Human somatic cell hybrid adherent macrovascular endothelial cell line, EA.hy926 (ATCC CRL^®^-2922^TM^) was used for the experiments. Ea.hy926 cells (ECs) were grown and maintained using Dulbecco’s Modified Eagle Medium (DMEM) with 10% FBS at 37 ℃. For the experiments, ECs of the passage S2 to S6 were used. The cell viability was determined using the Trypan Blue Exclusion assay (33).

### Preparation of drug solutions and treating with ECs

Three commercially available angiogenesis-promoting drugs, Phenytoin (Abbott India Ltd., Abbott Healthcare Pvt. Ltd., Mumbai, India), Vitamin C (Ascorbic acid, State Pharmaceuticals Manufacturing Corporation, No. 11, Sir John Kotelawala Mawatha, Ratmalana, Sri Lanka), and Simvastatin (Simvas-10; Micro Labs Limited, Sipcot, Hosur, India) were selected for the study. The solubility information for each drug was retrieved from the literature (13, 15, 34, 35). Phenytoin sodium and ascorbic acid were dissolved in complete culture media while Simvastatin was initially dissolved in Dimethyl Sulfoxide (DMSO), a cryopreserve agent. Since DMSO can damage cells at room temperature (RT), a highly concentrated Simvastatin solution was prepared using a minimum volume of DMSO and a dilution series was made using DMEM complete culture media (with 5% Bovine Serum Albumin; BSA). Since DMSO can damage cells at room temperature, the drug solutions were prepared so that the DMSO was highly diluted, with a dilution factor of at least 1:4000 in each solution. The dilution factor of DMSO in each concentration of the drug solution was calculated. All the drug solutions were prepared using DMEM culture media with 10% BSA (Bovine Serum Albumin). Sulforhodamine B (SRB) assay was conducted to test the cytotoxicity of each drug solution (36). Plasma concentrations of the drugs were obtained from the literature (13, 15, 34, 35) and a range of optimum concentrations for the SRB assay was determined for each drug as follows; The endothelial cells were plated in a 96-well cell culture plate at 5 x 10^4^ cells/well incubated overnight in a 37 ℃, 5% CO_2_ incubator. After confirming the attachment of the cells, drug solutions were added and incubated in the 5% CO_2_ incubator for 3 hours at 37 ℃ (Ascorbic acid 40, 50, 60, 70 and 80 µM; Phenytoin sodium 10, 15, 20, 25l, 30, and 40 µg/ml; Simvastatin 1, 0.5, 0.1, 0.05 and 0.01 µM). As controls, DMEM complete culture media and a dilution series of DMSO prepared using DMEM (diluted by 4000, 8000, 40,000, 80,000, 400,000-fold) were used. After the incubation, for each test, plate was washed twice with PBS and 40 µl of incomplete culture media at room temperature and 10 µl of Trichloroacetic acid was added to each well. The plate was then sealed with parafilm and was stored at 4 ℃ for 1 hour to fix the cells. The plate was washed five times and was air dried. 100 µl of SRB dye was added and incubated at room temperature for 30 minutes. The plate was then washed five times with glacial acetic acid. After air drying, 100 µl of Tris base was added and kept on a shaker for 1 hour. Absorbance was measured at 540nm. Using the SRB assay results, two concentrations for each drug were selected; the maximum non-toxic concentration and the minimum non-toxic concentration

### Assessment of cell-to-cell distances and cell morphology after treatment with sera and drug solutions

The cells were plated in a 96-well plate at a concentration of 7.5 x 10^4^ cells/well and incubated in a 5% CO_2_ incubator for 2 days at 37 ℃. When a cell monolayer was formed, DF, DHF (n= 5), HC (n= 5) sera, and the combinations of serum and drugs of selected concentrations were added in duplicates along with controls and incubated in a 5% CO2 incubator at 37℃ for 3 hours. This 3 hour incubation period was selected based on previous literature (37). After incubation, cells were stained with Giemsa working solution (1 ml of Giemsa stock solution dissolved in 9ml of autoclaved distilled water) and observed under x200 magnification using OLYMPUS CK40 inverted microscope and four fields per well were photographed. All the photos were taken using the same device at the same zoom level to obtain photos with equal resolution (4032×3024). The photos were brought to the same zoom level and same pixel value (1627px x 1627px) using Paint 3D software and cell distances were measured in pixel value using ImageJ software. Each treatment category consists of five age-, gender-, and residence-matched samples. For each sample, 100 distances were obtained. A total of 500 cell-to-cell distances were obtained for each treatment category. For the morphological analysis, a grid was used to randomly select 100 cells per treatment category to measure cell areas using ImageJ software. DF, DHF, and HC serum samples were diluted at a 1:1 ratio with incomplete culture media. The drug solutions were prepared using incomplete culture media (ICM) as the solvent. Cell counts for each treatment category were obtained by counting cells that appear in the squares of the grid. Each treatment category consisted of five samples and cell counts were taken from 100 squares per sample.

### Data analysis

Statistical analysis was performed using the IBM Statistical Package for the Social Sciences (SPSS) 20.0. A P value of <0.05 was considered significant, at 95% confidence interval. Kolmogorov-Smirnov (K-S) test was used to determine the distributions of variables. Descriptive statistics are given as mean ± SD. Comparison of single independent categorical variables with 2 groups was performed using Mann Whitney U test.

### Data availability

All relevant data are available in the article.

## Results

### Effect of DMSO on cell viability in cells treated with different concentrations of Simvastatin

Since simvastatin was prepared using DMSO, SRB assay was conducted to exclude the possible effect of DMSO itself in each dilution used by determining the cell viability. Table 1 shows the mean percentage cell viabilities obtained for DMSO dilutions. The cell viability observed with all DMSO dilutions (4000, 8000, 40,000, 80,000, and 400,000-fold) compared to the controls were comparable at 95% confidence level. Further, the percentage viabilities obtained for all DMSO dilutions were greater than 75%. Therefore, the effect of DMSO in the simvastatin preparations was considered negligible, not affecting the cell viability by itself, when compared to the controls.

**Table 1:** Mean percentage viability of cells for DMSO dilutions.

| DMSO dilution factor | Cell viability, Mean ± SD (%) | Significance (p-value) * |
| --- | --- | --- |
| 4000 | 79.15 ± 14.18 | 0.053 |
| 8000 | 79.36 ± 14.26 | 0.060 |
| 40,000 | 79.62 ± 14.27 | 0.066 |
| 80,000 | 79.88 ± 13.64 | 0.052 |
| 400,000 | 80.14 ± 13.54 | 0.054 |
| Control (with ICM) | 100.00 | - |

\*Significance between cells treated with DMSO and controls without treatment (p-value) were obtained using Kruskal-Wallis non-parametric test

#### Determination of drug concentrations for serum treatment assay

The cell viability for each concentration of the three drugs was compared with the cell viability of the controls after 3 hours of incubation. Table 2 shows the viability of cells treated with different concentrations of the three drugs.

**Table 2:**
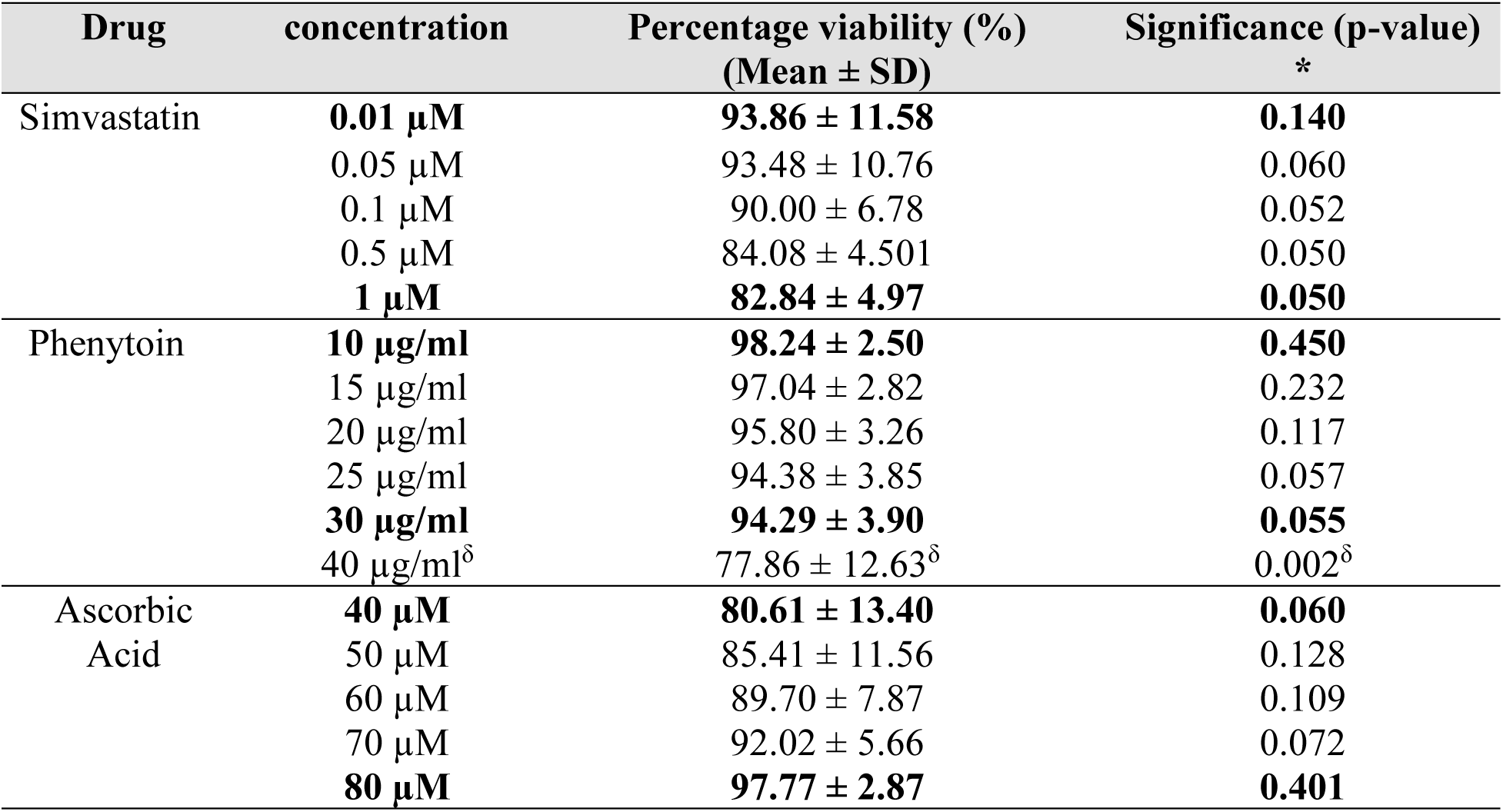
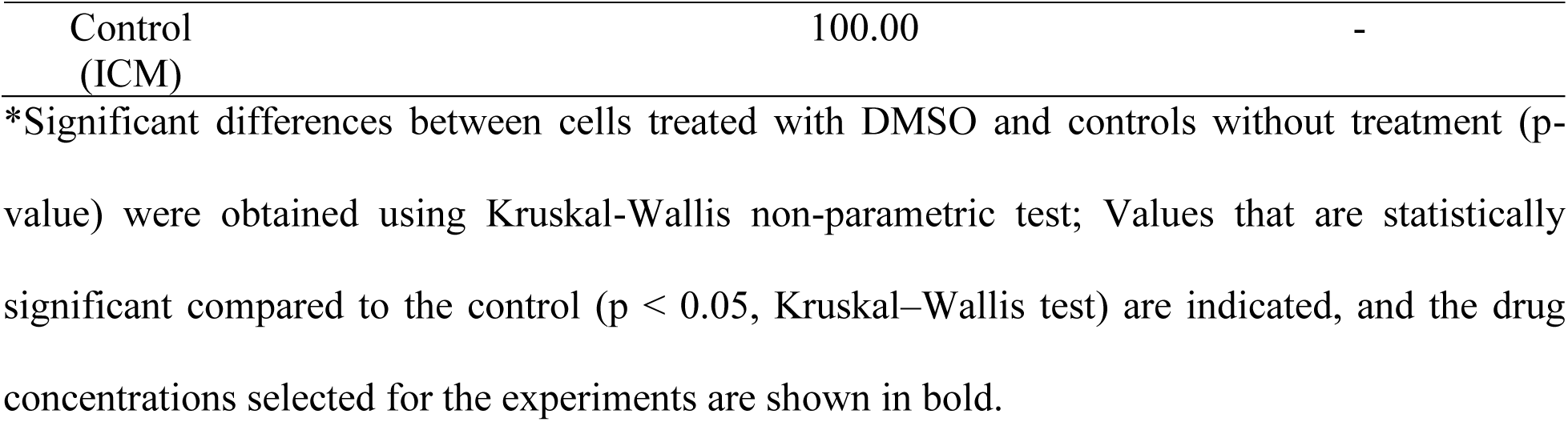
Viability of cells treated with different concentrations of simvastatin, phenytoin and ascorbic acid.

None of the five concentrations of the simvastatin showed a significant reduction in cell viability compared to the control at a 99.5% confidence level. Therefore, all concentrations of simvastatin tested were suitable to carry out the treatment experiments. The highest and the lowest non-toxic concentrations of simvastatin, 1 µM, and 0.01 µM respectively were selected to be used for downstream experiment.

All the concentrations of phenytoin, except 40 µg/ml, showed mean viability values of more than 80%. From the six concentrations tested, 40 µg/ml concentration showed a significant reduction in cell viability compared to the control at a 95% confidence level (p=0.020), indicating that the other concentrations (10, 15, 20, 25 and 30 µg/ml) of Phenytoin were suitable to carry out the treatment experiments. Therefore, the highest and the lowest nontoxic concentrations, 30 µg/ml and 10 µg/ml respectively, were selected for downstream experiments.

All the concentrations of ascorbic acid showed mean viabilities of more than 80% and showed no significant reduction in cell viability compared to the control at a 95% confidence level. Therefore, all concentrations of ascorbic acid tested (40, 50, 60, 70 and 80 µM) were suitable to carry out the experiments. The highest and the lowest nontoxic concentrations, 80 µM and 40 µM respectively, were selected.

#### Assessment of the changes in cell-to-cell distances and morphology of Ea.hy926 cells treated with DF, DHF, and HC sera and in combination with Simvastatin, Phenytoin and Ascorbic acid

##### Cell-to-cell distances of each treatment group

For each treatment category 500 cell-to-cell distances were measured from the ImageJ software using the photos of cells stained with Giemsa observed. Table 3 shows the average distances obtained for each treatment category after 3 hours of incubation.

**Table 3:** Average cell distances obtained i*n* Ea.hy926 cells after 3-hour incubation with each treatment category.

| Treatment category | Cell-to-cell distance of each treatment category (pixels) (Mean±SD) |
| --- | --- |
| Control group (cells + ICM with 5% BSA) | 7.459 ± 2.21 |
| Cells + Healthy control serum | 7.453 ± 1.53 |
| Cells + DF serum | 7.391 ± 1.64 |
| Cells + DHF serum | 16.903 ± 4.10 |
| Cells + DHF + Simvastatin 0.01 µM | 1.599 ± 3.53 |
| Cells + DHF + Simvastatin 1 µM | 6.044 ± 4.62 |
| Cells + DHF + Phenytoin 10 µg/ml | 23.734 ± 8.33 |
| Cells + DHF + Phenytoin 30 µg/ml | 23.308 ± 7.18 |
| Cells + DHF + Ascorbic acid 40 µM | 16.867 ± 6.71 |
| Cells + DHF + Ascorbic acid 80 µM | 16.784 ± 2.34 |

Figure 1 shows the mean cell-to-cell distances obtained for HC, DF, and DHF, in comparison to treatment with Simvastatin (0.01 µM and 1 µM), Ascorbic acid (40 µM and 80 µM), and Phenytoin (10 µg/ml and 30 µg/ml).

**Figure 1:**
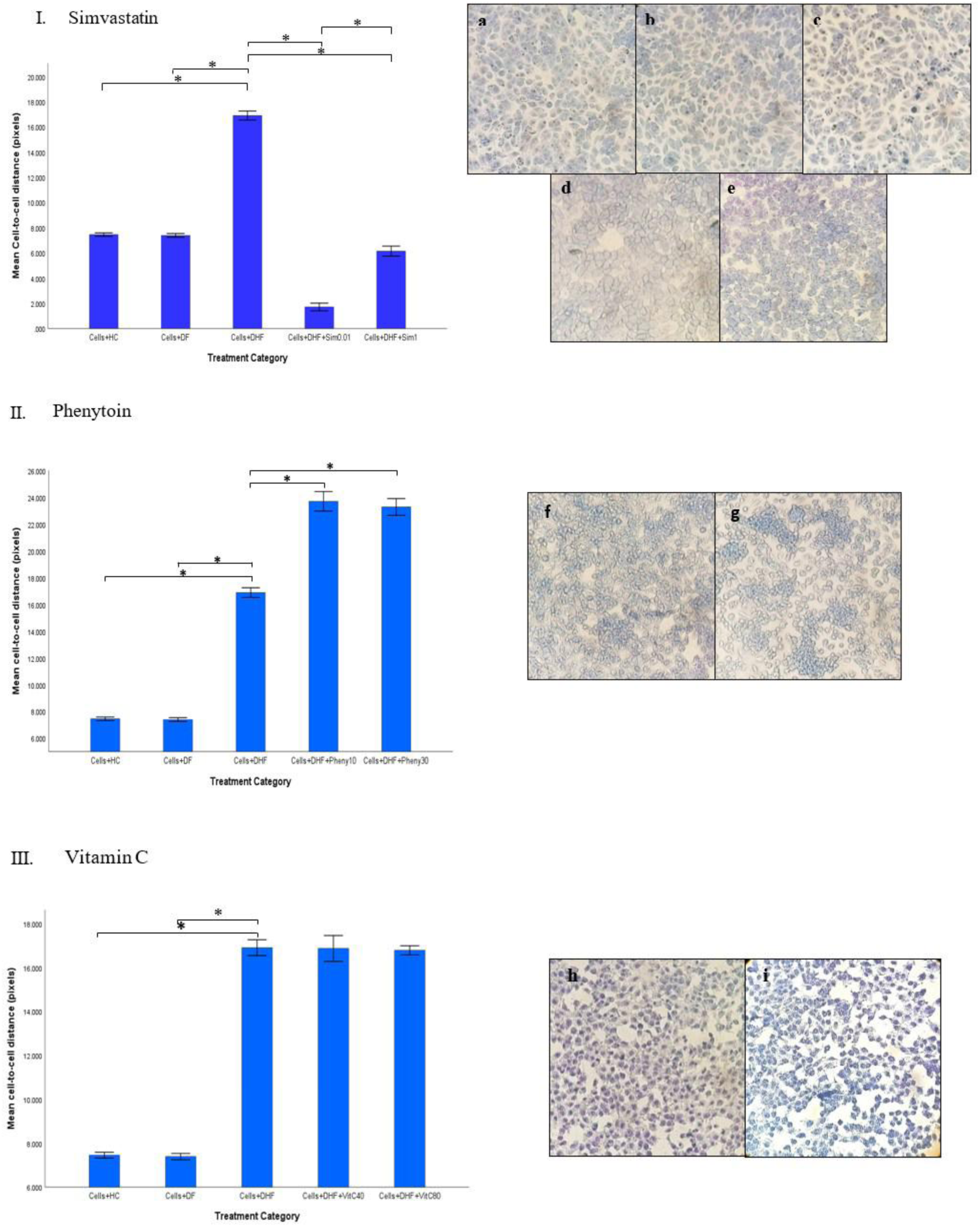
The comparison of cell distances obtained for different drug treatments. I) Simvastatin II) Phenytoin III) Vitamin C and the appearance of Giemsa-stained cells after 3 hours of treatment with I-a) HC serum, I-b) DF serum, I-c) DHF serum, I-d) 0.01 µM Simvastatin+DHF serum, I-e) 1 µM Simvastatin+DHF serum, II-f) 10 µM Phenytoin+DHF serum, II-g) 30 µM Phenytoin+DHF serum, III-h) 40 µM Vitamin C+DHF serum, III-) 80 µM Vitamin C+DHF serum (x200) No significant difference in cell distances was observed between the cells treated with HC serum and DF serum (p= 0.435). However, cells treated with DHF serum showed a significant increase in cell-to-cell distances compared to cells treated with HC serum (p<0.05) and cells treated with DF serum (p<0.05), demonstrating endothelial monolayer destabilization that occurs in the presence of DHF serum (Figure 1(I-c)).

Cells treated with a combination of DHF serum and Simvastatin (0.01 µM and 1 µM) showed a significant reduction in the cell-to-cell distances compared to cells treated with DHF serum alone (p<0.05), indicating a reduction of cell-to-cell gaps after Simvastatin treatment. Between the two concentrations of Simvastatin, a significant reduction of cell-to-cell distance was observed for 0.01 µM, compared to 1 µM, in the presence of DHF serum (p<0.05).

No significant difference in distances between cells was observed for cells treated with Ascorbic acid (40 µM and 80 µM) in the presence of DHF serum compared to cells treated with DHF serum alone (Average cell-to-cell distance 40 µM ascorbic acid + DHF serum =16.867 ± 6.708, Average cell-to-cell distance 80 µM ascorbic acid + DHF serum =16.784 ± 2.343, Average cell-to-cell distance DHF serum = 16.903 ± 4.096; p=0.059, p=0.436). However, a significant increase in cell distances was observed in cells treated with Phenytoin (10 µg/ml and 30 µg/ml) combined with DHF serum compared to cells treated only with DHF serum (p<0.05).

##### Morphological analysis

After Giemsa staining, the cells of each treatment category were observed for morphological changes between the categories. The EC shape varies along the vascular tree. The ECs treated with DHF sera combined with different drugs were observed to have morphologies that deviated from the commonly considered thin, slightly elongated spindle-shape with smooth, regular cell margins. Figure 1 shows the images of Giemsa-stained cells treated with HC serum, DHF serum, DHF combined with ascorbic acid, DHF combined with phenytoin, and DHF combined with Simvastatin. The cells treated with HC serum (Figure 1(I-a)) and DF serum (Figure 1(I-b)) are situated more closely to one another compared to cells treated with DHF serum (Figure 1(I-c)). Further, the cells in Figure 1(I-a) are spindle-shaped with smooth regular cell membranes. Interestingly, the cells treated with DHF serum combined with both 0.01 µM and 1 µM Simvastatin (Figures 1(I-d) & 1(I-e)) have lost the spindle shape but show the formation of a tight monolayer with cells being tightly joined to the adjacent cells. The cells treated with DHF serum combined with both concentrations of Phenytoin (Figures 1(II-f) & 1(II-g) appear to have shrunken in size with irregular cell shapes and noticeable cell damage. The cells treated with DHF serum combined with both concentrations of Ascorbic acid appear to have irregular cell membranes (Figures 1(III-h) & 1(III-i)). The spindle shape of the ECs is not observed here.

Simvastatin-treated cells showed a significant reduction in the cell-to-cell distances compared to DHF serum-treated cells, especially at 0.01 µM (p<0.05). Further, a noticeable change in the cell morphology was also observed. Therefore, the cell areas of 0.01 µM and 1 µM Simvastatin treatment categories were measured using ImageJ software and compared against the cells of HC, DF, and DHF treatment categories to see if treatment with Simvastatin has caused a significant difference in the cell size (cell area). Since Ascorbic acid showed no significant effect on cell-to-cell distances compared to DHF-treated cells, and Phenytoin showed endothelial cell damage when combined with DHF sera, cell size analysis was done only for Simvastatin.

ImageJ software was used to measure areas of 20 cells per sample for each of the HC, DF, DHF and DHF combined with 0.01 µM and 1 µM Simvastatin treatment categories. The areas were obtained in square pixels. Since all the photos are of the same zoom level and same pixel value (1627×1627), comparisons can be made between the measurements obtained.

Table 4 shows the average area of cells for each treatment category and Figure 2 shows a comparison of average cell areas between HC, DF, DHF, 0.01µM Simvastatin and 1µM Simvastatin + DHF serum treatment categories.

**Table 4:** Area of cell in each treatment categories.

| Treatment category | Average Area of 100 cells (pixel<sup>2</sup>) |
| --- | --- |
| Cells + HC serum | 4564.1 ± 1528.2 |
| Cells + DF serum | 4429.0 ± 1305.9 |
| Cells + DHF serum | 4397.1 ± 1489.1 |
| **Cells + DHF + Simvastatin 0.01 μM** | **7092.2 ± 2120.7** |
| Cells + DHF + Simvastatin 1 μM | 4953.2 ± 1885.2 |

Significant increase in cell size in cells exposed to 0.01 μM Simvastatin compared to other categories including DHF serum+1 μM Simvastatin (p<0.05) is given in bold font.

**Figure 2:**
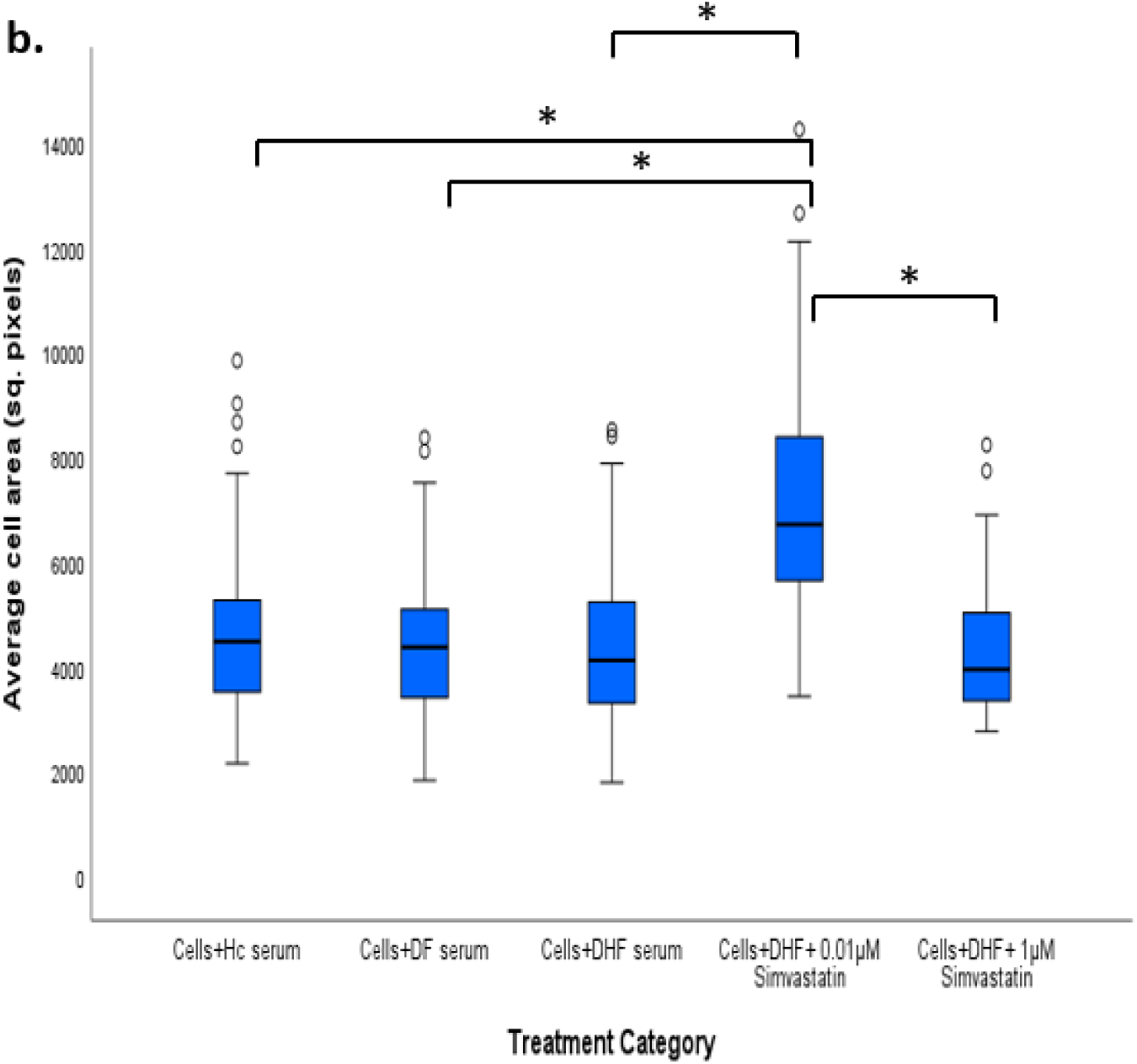
Comparison of average cell areas between HC, DF, DHF, 0.01µM Simvastatin and 1µM Simvastatin + DHF serum treatment categories (*Significant difference in cell areas among different categories) The cells treated with HC serum, DF serum, and DHF serum showed no significant difference in average cell areas, when compared against each other (p>0.05). The results depicted a significant increase in cell size in cells exposed to 0.01 µM Simvastatin compared to other categories including DHF serum+1µM Simvastatin (p<0.05).

Table 5 shows the average cell counts obtained for each treatment category and Figure 3 shows a comparison of the cell counts obtained for HC, DF, DHF, 0.01µM Simvastatin and 1µM Simvastatin + DHF serum treatment categories. Results showed no significant difference in the cell counts between the categories (p>0.05).

**Table 5:** Average cell counts obtained for each treatment categories.

| Treatment category | Average cell count |
| --- | --- |
| Cells + HC serum | $17.71 \pm 3.19$ 365 |
| Cells + DF serum | $17.72 \pm 2.85$ |
| Cells + DHF serum | $15.95 \pm 2.98$ 366 |
| Cells + DHF + Simvastatin 0.01 μM | $17.02 \pm 3.19$ |
| Cells + DHF + Simvastatin 1 μM | $17.68 \pm 4.53$ 367 |

**Figure 3:**
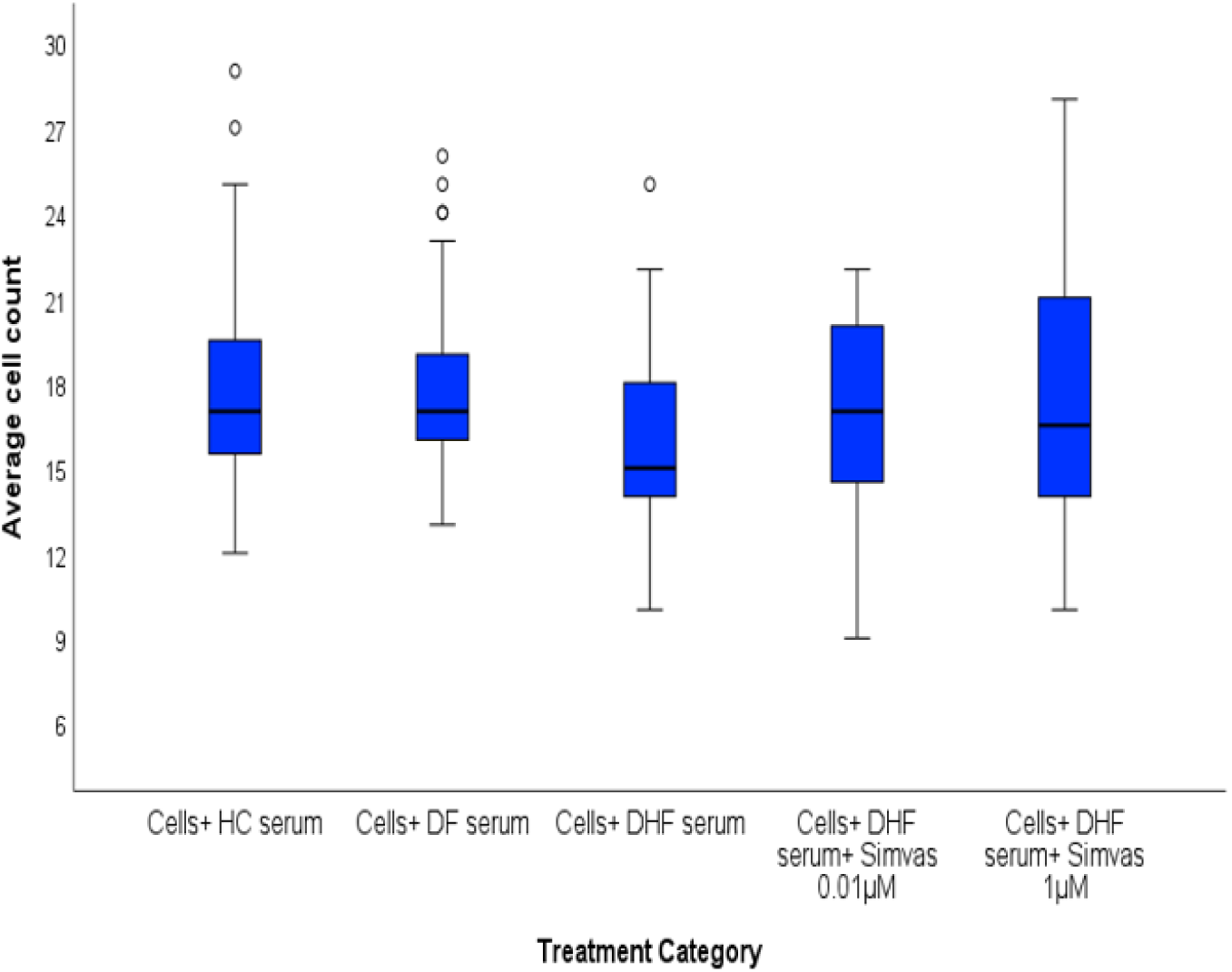
Comparison of average cell counts obtained for each treatment category.

## Discussion

In this study, we focused on three commercially available drugs that have pleiotropic effects on angiogenesis promotion, on their ability *in vitro* to inhibit the endothelial monolayer destabilization caused in the presence of DHF serum. We have demonstrated that the endothelial monolayer disintegrates when treated with DHF sera and that in the presence of 0.01 µM and 1 µM Simvastatin, this disintegration could be inhibited/reversed, resulting in interdigitation of the ECs into a monolayer. Further, it was observed that the Simvastatin treatment in the presence of DHF serum resulted in a monolayer morphologically similar to a cobblestone monolayer. It was also observed that treatment with Ascorbic acid does not result in a significant change in the monolayer destabilization caused by the DHF sera, while Phenytoin showed evidence of promoting the destabilization.

When the endothelial monolayer was treated with DHF sera, the average cell-to-cell distances between the adjacent cells increased in comparison to cells treated with HC sera and cells treated with DF sera (DHF serum treatment = 16.90 ± 4.09; p<0.05, HC serum treatment = 7.45 ± 1.53, DF serum treatment = 7.39 ± 1.64) at 95% confidence level. This demonstrates that in the presence of DHF sera, the endothelial monolayer destabilizes *in vitro*. This *in vitro* observation indicates to similar effects of DHF sera *in vivo* which may lead to increased permeability in DHF patients. Although the exact mechanism behind the observed results has not been studied here, it could be supported by several previous studies which had shown increased endothelial permeability due to changes in expression of cytoadhesion molecules. A study conducted by Luplerdlop et al. in 2006 found that primary human umbilical vascular endothelial cell (HUVEC) monolayers treated with supernatant of immature dendritic cell (iDC) cultures infected with DV, resulted in increased permeability of the endothelium, mainly due to the loss of expression of cell adhesion molecules (38). The expression of platelet endothelial adhesion molecule 1 (PECAM-1) and vascular endothelium (VE)-cadherin (cell adhesion molecules) were found to have decreased significantly. Further, it was discovered that the decrease in the expression of cell adhesion molecules is associated with increased levels of matrix metalloproteases (MMPs) resulting in increased endothelial permeability, affecting the monolayer formation (39).

MMPs are zinc-dependent endopeptidases that enhance endothelial permeability. They are produced by ECs in response to pro-inflammatory cytokines such as TNF-α (40). Pro-inflammatory cytokines like TNF-α are found to be at elevated levels in dengue patients according to a study conducted by (41). In a study conducted by Luplerdlop et al. in 2006, the dengue virus-infected immature dendritic cells were found to overproduce (MMP)-9 in large quantities while also producing MMP-2 in smaller quantities. In another study conducted by Zuo et al. in 2014, it was found that MMP is upregulated in macrophages upon DENV infection (42). When present in excess, the MMPs are found to have deleterious effects on EC integrity. Therefore, it is possible that the serum obtained from the DHF patients contains MMPs in elevated levels in addition to the elevated Ang-2 levels. Elevation of MMPs however was not tested and confirmed in the current study.

According to the results obtained, significant reductions in cell-to-cell distances were observed in the cells treated with 0.01 µM (Figure 1-a) and 1 µM (Figure 1-b) Simvastatin when combined with DHF serum, compared to the cells treated with just the DHF serum (Figure 1-c) (a, 1.60 ± 3.53; b, 6.04 ± 4.62; c, 16.90 ± 4.10; p<0.050). Both 1 µM and 0.01 µM concentrations of Simvastatin showed a significant reduction in cell-to-cell distances in the presence of DHF sera. When compared to each other, low-dose (0.01 µM) Simvastatin showed the highest reduction in cell distances among the two. Therefore, it is noteworthy that the low-dose Simvastatin (0.01 µM) showed a significantly higher reduction in the cell-to-cell distances in the presence of DHF serum, possibly demonstrating the formation of EC monolayer even in the presence of serum factors leading to EC monolayer destabilization. However, it should be noted that 1 µM Simvastatin, while reducing the cell-to-cell distances of cells in the presence of DHF serum, restores the cell distances to values similar to cells treated with HC serum. Therefore, it can be argued that 1 µM dose of Simvastatin could be a better candidate for future experiments.

One explanation behind the ability of Simvastatin to promote the formation of a monolayer could be attributed to its ability to lower the Ang-2 levels. In a study conducted by Tuuminen et al. in 2014, Simvastatin was associated with lowering intravitreal Ang-2 levels. The study has found that diabetic patients with macular edema (DME) and patients with proliferative diabetic retinopathy (PDR) who received simvastatin treatment showed reduced levels of Ang-2 compared to patients without statin medication (43). Angiopoietin-2 (Ang-2) has been shown to induce endothelial destabilization resulting in plasma leakage that is characteristic of severe dengue disease, DHF. Further, it was recently observed that Ang-2 in human blood rises when a person approaches a leaking phase (critical stage of the disease) of severe dengue indicating Ang-2 involvement in inducing DHF possibly through endothelial dysfunction (7, 44).

Simvastatin also can inhibit the production of MMP-9 in human ECs. Many studies have shown a role for several members of the statin family in suppressing the expression of MMPs in macrophages and ECs. In a study conducted by Massaro et al. in 2010, it was shown that human umbilical vein ECs (HUVEC) treated 0with Simvastatin inhibited the matrix metalloproteinase-9 (45). As mentioned earlier, the increased permeability of the endothelium, resulting from the elevated levels of MMPs, was discovered to be associated with a loss of expression of the PECAM-1 and VE-cadherin cell adhesion molecules (42). The inhibition of the production of MMP-9 by Simvastatin could have resulted in increased expression of EC adhesion molecules such as PECAM-1 and VE-cadherin. This could be another reason behind ECs showing tight cell-cell contacts with the adjoining cells after the treatment with Simvastatin in the current study. Therefore, it is suggestive that Simvastatin may stabilize the endothelium by reducing the serum Ang-2 levels or by inhibiting the MMP-9 in human ECs and these suggestions have yet to be clarified in future experiments.

According to the results, both concentrations of ascorbic acid (40 µM and 80 µM) showed no significant change in cell-to-cell distances in the presence of DHF serum, indicating that ascorbic acid had no significant effect, positive or negative, towards the endothelial monolayer destabilization in the presence of DHF serum. However, both concentrations of phenytoin (10 µM and 30 µM) showed a significant increase in the average cell-to-cell distances compared to cells treated with DHF. Although not many studies have been conducted to test the effects of Phenytoin on vascular ECs, a recent experimental in vitro study by Ballesteros-Peña, S et al. in 2022, identified phenytoin to be an alkaline that shows extreme osmolarity values in undiluted form and maintains high tonicity values after dilution in 100ml saline. Accordingly, since irritation and endothelial damage could be promoted by fluids with high osmolality and alkaline pH, Phenytoin could have the capacity to cause endothelial damage and irritation when administered intravenously (46). However, further studies need to be done to explain the observations obtained in this study. Since Phenytoin is a commonly used anti-convulsant drug to treat dengue patients undergoing seizures due to encephalopathy, it is important that the combinatorial effects of Phenytoin in the presence of DHF serum factors to be tested thoroughly in further studies.

The cells treated with both concentrations of Simvastatin showed an increase in the average cell size. Cells treated with 0.01 µM Simvastatin showed a significant increase compared to cells treated with HC, DF and DHF serum (p<0.05). Although cells treated with 1 µM Simvastatin showed an increase in the average cell size compared to HC, DF and DHF serum treated cells, the change was not significant (p>0.05). However, cells treated with 0.01 µM Simvastatin showed a significant increase in the cells size compared to cells treated with 1 µM Simvastatin. A study conducted by Miettinen & Björklund in 2015 found that low concentrations of statin drugs limit proliferation and thus increase cell size. Higher concentrations of the statins could limit cell growth possibly through inhibiting Mammalian target of rapamycin (mTOR) and other growth promoting signals or by having off target effects that block growth (47). This may partly explain why 0.01 µM Simvastatin treatment showed a significant increase in the average cell size compared to the other treatment categories while 1 µM Simvastatin treatment resulted in an increase in cell size, but to a lesser extent which is statistically insignificant.

ECs in the monolayer treated with DHF serum increases in the cell-to-cell distances without a change in the cell size or the cell count compared to HC or DF serum treated cells. However, it is noteworthy that cells treated with Simvastatin 0.01 µM in the presence of DHF reduces cell-cell distances while showing a significant increase in the cell size and no significant change in the cell count compared to HC and DF serum treated cells. This implies that reduction in cell-to-cell distances of cells treated with low-dose Simvastatin is not due to cell proliferation, but could mainly be due to the increase in the average cell size, which is a known effect of Simvastatin. This eliminates the assumption that Simvastatin might have reduced the cell-to-cell distances through cell proliferation, which could have implications in risk of cancer.

Another interesting observation was made regarding the morphology of the ECs after the drug treatments. The morphology of the cells treated with DHF serum combined with 0.01 µM and 1 µM Simvastatin showed a deviation from the spindle shape of the untreated endothelial cells or cells treated with HC serum. The morphology also appears to be similar to the cells treated with only DHF serum. In the presence of Simvastatin, the cells appeared to take a round shape with regular cell membranes, with no damaged or lysed cells. Most importantly, cells treated with both concentrations of Simvastatin (0.01 µM and 1 µM) showed interdigitating at EC borders through typical zigzag lines, evidencing the formation of EC monolayer, with cells being tightly joined to the adjacent cells. Observing the images of HUVEC monolayers generated in a study by Jiménez N et al in 2013, which shows similar morphology, it is possible that the Simvastatin treatment, in the presence of DHF, could have promoted the establishment of cobblestone monolayer (48, 49). The exact reason behind this formation of the cobblestone monolayer in ECs treated with Simvastatin in the presence of DHF has not yet been studied. TNF-α is thought to promote the atheroprone morphology in ECs and is used as an inflammatory cytokine in adhesion experiments (49). ECs simulated by TNF-α have been found to assume an inflammatory phenotype with elevated levels of atheroprone markers such as ICAM-1 and VCAM-1 (50, 51). In a study conducted by Dick M et al. in 2015, it was found that although statins are known to be beneficial to ECs and promote atheroprotective genotype, they influence the ECs to become rounder under specific conditions (52). Since DHF serum is found to be rich in TNF-α, and statins have been found to influence the ECs to become rounder, the formation of the cobblestone monolayer could be a combinatorial effect of these two factors. With the light shed by this study, further experiments could be designed to test such combinatorial effects of the statin drugs and the serum factors in DHF patients, to develop an effective therapy to treat DHF. Further experiments should be done to explore the effects of Simvastatin on the morphology of vascular ECs using fluorescent staining methods. The pathways and mechanisms underlying the promotion of the monolayer formation and its stabilization should also be further investigated.

## Conclusion

This was a preliminary study which was conducted to test whether Simvastatin, Phenytoin and Vitamin C could reverse/inhibit the endothelial monolayer destabilization caused in the presence of DHF serum, *in vitro*. Simvastatin treatment in the presence of DHF serum showed significant reduction in cell-to-cell distances and monolayer formation in Ea.hy926 cells for both 1 µM and 0.01 µM concentrations. Further, Simvastatin treatment resulted in an increase in cell sizes for both 0.01 µM and 1 µM concentrations, among which 0.01 µM concentration showed a statistically significant increase. No significant change was observed in the cell counts for cells treated Simvastatin compared to HC, DF and DHF treatment categories. Phenytoin showed a significant increase in the cell-to-cell distances and cell damage in the presence of DHF serum while Ascorbic acid showed neither a positive, nor a negative effect on the endothelial monolayer formation. This work suggests that low dose Simvastatin could have a role in promoting endothelial monolayer formation and its stabilization even in the presence of DHF serum and that the reduction in cell-to cell distances is not due to cell proliferation, but could mainly be due to the increase in the average cell size, which is a known effect of Simvastatin. Further studies are needed to explore the pathways and mechanisms by which Simvastatin could promote the endothelial monolayer formation.

## Acknowledgements

Assistance rendered by the Institute of Biochemistry, Molecular Biology and Biotechnology, Department of Zoology and Environment Sciences, Faculty of Science, University of Colombo, Sri Lanka, North Colombo Teaching Hospital, Ragama, Sri Lanka, and the financial assistance from the University of Colombo are acknowledged. We also wish to thank all the hospital staff, patients, their family members and healthy individuals for their voluntary participation in this study.

## References

1. Bhatt S, Gething PW, Brady OJ, Messina JP, Farlow AW, Moyes CL, Drake JM, Brownstein JS, Hoen AG, Sankoh O, Myers MF, George DB, Jaenisch T, Wint GRW, Simmons CP, Scott TW, Farrar JJ, Hay SI. 2013. The global distribution and burden of dengue. Nature 496:504–507.

2. Tissera H, Ooi E, Gubler D, Tan Y, Logendra B, Wahala W, De Silva A, Abeysinghe MRN, Palihawadana P, Gunasena S, Tam C, Amarasinghe A, Letson GW, Margolis H, De Silva A. 2011. New Dengue Virus Type 1 Genotype in Colombo, Sri Lanka. Emerg Infect Dis 17.

3. Ali S, Khan AW, Taylor-Robinson AW, Adnan M, Malik S, Gul S. 2018. The unprecedented magnitude of the 2017 dengue outbreak in Sri Lanka provides lessons for future mosquito-borne infection control and prevention. Infect Dis Health 23:114–120.

4. Halstead S. 2019. Handbook of Viral and Rickettsial Hemorrhagic Fevers. CRC Press.

5. Martina BEE, Koraka P, Osterhaus ADME. 2009. Dengue Virus Pathogenesis: an Integrated View. Clin Microbiol Rev 22:564–581.

6. Michels M, Van Der Ven AJAM, Djamiatun K, Fijnheer R, De Groot PG, Griffioen AW, Sebastian S, Faradz SMH, De Mast Q. 2012. Imbalance of Angiopoietin-1 and Angiopoetin-2 in Severe Dengue and Relationship with Thrombocytopenia, Endothelial Activation, and Vascular Stability. Am Soc Trop Med Hyg 87:943–946.

7. Mapalagamage M, Handunnetti SM, Wickremasinghe AR, Premawansa G, Thillainathan S, Fernando T, Kanapathippillai K, De Silva AD, Premawansa S. 2020. High Levels of Serum Angiopoietin 2 and Angiopoietin 2/1 Ratio at the Critical Stage of Dengue Hemorrhagic Fever in Patients and Association with Clinical and Biochemical Parameters. J Clin Microbiol 58:e00436–19.

8. Korhonen EA, Lampinen A, Giri H, Anisimov A, Kim M, Allen B, Fang S, D’Amico G, Sipilä TJ, Lohela M, Strandin T, Vaheri A, Ylä-Herttuala S, Koh GY, McDonald DM, Alitalo K, Saharinen P. 2016. Tie1 controls angiopoietin function in vascular remodeling and inflammation. J Clin Invest 126:3495–3510.

9. Luplerdlop N, Missé D, Bray D, Deleuze V, Gonzalez J, Leardkamolkarn V, Yssel H, Veas F. 2006. Dengue-virus-infected dendritic cells trigger vascular leakage through metalloproteinase overproduction. EMBO Rep 7:1176–1181.

10. Zuo X, Pan W, Feng T, Shi X, Dai J. 2014. Matrix Metalloproteinase 3 Promotes Cellular Anti-Dengue Virus Response via Interaction with Transcription Factor NFκB in Cell Nucleus. PLoS ONE 9:e84748.

11. Malavige G, Ogg G. 2024. Molecular mechanisms in the pathogenesis of dengue infections. Trends in Molecular Medicine 30:484–498.

12. Wu H, Jiang H, Lu D, Qu C, Xiong Y, Zhou D, Chopp M, Mahmood A. 2011. Induction of Angiogenesis and Modulation of Vascular Endothelial Growth Factor Receptor-2 by Simvastatin After Traumatic Brain Injury. Neurosurgery 68:1363–1371.

13. Medina RJ, O’Neill CL, Devine AB, Gardiner TA, Stitt AW. 2008. The Pleiotropic Effects of Simvastatin on Retinal Microvascular Endothelium Has Important Implications for Ischaemic Retinopathies. PLoS ONE 3:e2584.

14. Chade AR, Zhu X, Mushin OP, Napoli C, Lerman A, Lerman LO, Chade AR, Zhu X, Mushin OP, Napoli C, Lerman A, Lerman LO. 2006. Simvastatin promotes angiogenesis and prevents microvascular remodeling in chronic renal ischemia. FASEB J 20:1706–1708.

15. Duarte JA, Barros ALB de, Leite EA. 2021. The potential use of simvastatin for cancer treatment: A review.

16. Stine JE, Schointuch M, Zhou C, Gilliam T, Han X, Gehrig PA, Bae-Jump VL. 2014. Simvastatin, an HMG-CoA reductase inhibitor, exhibits antimetastatic and antitumorigenic effects in endometrial cancer 133:2–207.

17. Tatè R, Zona E, Cicco Rosanna De, Trotta V, Urciuoli M, Morelli A, Baiano S, Carnuccio R, Fuggetta MP, Morelli F. 2017. Simvastatin inhibits the expression of stemness-related genes and the metastatic invasion of human cancer cells via destruction of the cytoskeleton 51:1851–1859.

18. Bersudsky Y. 2006. Phenytoin: an anti-bipolar anticonvulsant? 9:479–484.

19. Anstead GM, Hart LM, Sunahara JF, Liter ME. 1996. Phenytoin in Wound Healing. Ann Pharmacother 30:768–775.

20. Shaw J, Hughes CM, Lagan KM, Bell PM. 2007. The clinical effect of topical phenytoin on wound healing: a systematic review. Br J Dermatol 157:997–1004.

21. Wilson EL, Garton M, Fuller HR. 2016. Anti-epileptic drugs and bone loss: Phenytoin reduces pro-collagen I and alters the electrophoretic mobility of osteonectin in cultured bone cells. Epilepsy Res 122:97–101.

22. Corrêa JD, Queiroz-Junior CM, Costa JE, Teixeira AL, Silva TA. 2011. Phenytoin-Induced Gingival Overgrowth: A Review of the Molecular, Immune, and Inflammatory Features. ISRN Dent 2011:1– 8.

23. Turan M, Saraydyn SU, Bulut HE, Elagöz S, Cetinkaya O, Karadayi K, Canbay E, Sen M. 2004. Do vascular endothelial growth factor and basic fibroblast growth factor promote phenytoin’s wound healing effect in rat? An immunohistochemical and histopathologic study. Dermatol Surg 30:1303– 1309.

24. DaCosta ML, Regan MC, Al Sader M, Leader M, Bouchier-Hayes D. 1998. Diphenylhydantoin sodium promotes early and marked angiogenes is and results in increased collagen deposition and tensile strength in healing wounds. Surgery 123:287–293.

25. Pitiakoudis M, Giatromanolaki A, Iliopoulos I, Tsaroucha A, Simopoulos C, Piperidou C. 2004. Phenytoin-Induced Lymphocytic Chemotaxis, Angiogenesis and Accelerated Healing of Decubitus Ulcer in a Patient with Stroke. J Int Med Res 32:201–205.

26. May JM, Harrison FE. 2013. Role of Vitamin C in the Function of the Vascular Endothelium. Antioxid Redox Signal 19:2068–2083.

27. Telang S, Clem AL, Eaton JW, Chesney J. 2007. Depletion of Ascorbic Acid Restricts Angiogenesis and Retards Tumor Growth in a Mouse Model. Neoplasia 9:47–56.

28. May JM, Qu Z. 2005. Transport and intracellular accumulation of vitamin C in endothelial cells: relevance to collagen synthesis. Arch Biochem Biophys 434:178–186.

29. Vissers MC, Dias AB. 2020. Ascorbate as an enzyme cofactor, p. 71–98. *In* In Vitamin C. CRC Press.

30. Daghini E, Zhu X-Y, Versari D, Bentley MD, Napoli C, Lerman A, Lerman LO. 2007. Antioxidant vitamins induce angiogenesis in the normal pig kidney. Am J Physiol-Ren Physiol 293:F371–F381.

31. Levine GN, Frei B, Koulouris SN, Gerhard MD, Keaney JF, Vita JA. 1996. Ascorbic acid reverses endothelial vasomotor dysfunction in patients with coronary artery disease. Circulation 10.1161/01.CIR.93.6.1107.

32. Levine GN, Frei B, Koulouris SN, Gerhard MD, Keaney JF, Vita JA. 1996. Ascorbic Acid Reverses Endothelial Vasomotor Dysfunction in Patients With Coronary Artery Disease. Circulation 93:1107– 1113.

33. Strober W. 1997. Trypan Blue Exclusion Test of Cell Viability. Curr Protoc Immunol 21.

34. Frikke-Schmidt H, Lykkesfeldt J. 2010. Keeping the intracellular vitamin C at a physiologically relevant level in endothelial cell culture. Anal Biochem 397:135–137.

35. Patocka J, Wu Q, Nepovimova E, Kuca K. 2020. Phenytoin – An anti-seizure drug: Overview of its chemistry, pharmacology and toxicology. Food Chem Toxicol 142:111393.

36. Vichai V, Kirtikara K. 2006. Sulforhodamine B colorimetric assay for cytotoxicity screening. Nat Protoc 1:1112–1116.

37. Scharpfenecker M, Fiedler U, Reiss Y, Augustin HG. 2005. The Tie-2 ligand Angiopoietin-2 destabilizes quiescent endothelium through an internal autocrine loop mechanism. J Cell Sci 118:771– 780.

38. Luplerdlop N, Missé D, Bray D, Deleuze V, Gonzalez J, Leardkamolkarn V, Yssel H, Veas F. 2006. Dengue-virus-infected dendritic cells trigger vascular leakage through metalloproteinase overproduction. EMBO Rep 7:1176–1181.

39. Zuo X, Pan W, Feng T, Shi X, Dai J. 2014. Matrix Metalloproteinase 3 Promotes Cellular Anti-Dengue Virus Response via Interaction with Transcription Factor NFκB in Cell Nucleus. PLoS ONE 9:e84748.

40. Hummel V, Kallmann BA, Wagner S, Füller T, Bayas A, Tonn JC, Benveniste EN, Toyka KV, Rieckmann P. 2001. Production of MMPs in Human Cerebral Endothelial Cells and Their Role in Shedding Adhesion Molecules. J Neuropathol Exp Neurol 60:320–327.

41. Masood KI, Jamil B, Rahim M, Islam M, Farhan M, Hasan Z. 2018. Role of TNF α, IL-6 and CXCL10 in Dengue disease severity. Iran J Microbiol 10:202–207.

42. Luplerdlop N, Missé D, Bray D, Deleuze V, Gonzalez J, Leardkamolkarn V, Yssel H, Veas F. 2006. Dengue-virus-infected dendritic cells trigger vascular leakage through metalloproteinase overproduction. EMBO Rep 7:1176–1181.

43. Tuuminen R, Sahanne S, Loukovaara S. 2014. Low intravitreal angiopoietin-2 and VEGF levels in vitrectomized diabetic patients with simvastatin treatment. Acta Ophthalmol (Copenh) 92:675–681.

44. Rothman AL, Ennis FA. 1999. Immunopathogenesis of Dengue Hemorrhagic Fever. Virology 257:1– 6.

45. Massaro M, Zampolli A, Scoditti E, Carluccio MA, Storelli C, Distante A, De Caterina R. 2010. Statins inhibit cyclooxygenase-2 and matrix metalloproteinase-9 in human endothelial cells: anti-angiogenic actions possibly contributing to plaque stability. Cardiovasc Res 86:311–320.

46. Ballesteros-Peña S, Fernández-Aedo I, Vallejo-De La Hoz G, Tønnesen J, Miguelez C. 2022. Identification of potentially irritating intravenous medications. Enferm Intensiva Engl Ed 33:132– 140.

47. Miettinen TP, Björklund M. 2015. Mevalonate Pathway Regulates Cell Size Homeostasis and Proteostasis through Autophagy. Cell Rep 13:2610–2620.

48. Jiménez N, Krouwer VJD, Post JA. 2013. A new, rapid and reproducible method to obtain high quality endothelium in vitro. Cytotechnology 65:1–14.

49. Rouleau L, Copland IB, Tardif J-C, Mongrain R, Leask RL. 2010. Neutrophil Adhesion on Endothelial Cells in a Novel Asymmetric Stenosis Model: Effect of Wall Shear Stress Gradients. Ann Biomed Eng 38:2791–2804.

50. Bergh N, Ulfhammer E, Glise K, Jern S, Karlsson L. 2009. Influence of TNF-α and biomechanical stress on endothelial anti- and prothrombotic genes. Biochem Biophys Res Commun 385:314–318.

51. Rossi J, Rouleau L, Emmott A, Tardif J-C, Leask RL. 2010. Laminar shear stress prevents simvastatin-induced adhesion molecule expression in cytokine activated endothelial cells. Eur J Pharmacol 649:268–276.

52. Dick M, MacDonald K, Tardif J-C, Leask RL. 2015. The effect of simvastatin treatment on endothelial cell response to shear stress and tumor necrosis factor alpha stimulation. Biomed Eng OnLine 14:58.

